# VIVA (VIsualization of VAriants): A VCF file visualization tool

**DOI:** 10.1101/589879

**Authors:** George A. Tollefson, Jessica Schuster, Fernando Gelin, Ashok Ragavendran, Isabel Restrepo, Paul Stey, James Padbury, Alper Uzun

## Abstract

The volume and pace of data accumulation from high-throughput sequencing studies have been amplified by recent rapid technological advances in biological sciences. Visualization of genomic data is essential for quality control, exploration, and interpretation. Here, we describe a user-friendly visualization tool for variant call format (VCF) files with which the users can interactively evaluate and share genomic data, as well as create publication quality graphics.

## 1. INTRODUCTION

Next generation sequencing produces an enormous amount of genomic data. The volume of the genomic information varies based on the study. Many different types of file formats are generated during the variant discovery process. One file format commonly used in sequence analysis is the variant call format (VCF). It is a text file format generated during the variant calling process that contains information about variant positions in the genome. The structure includes variant information such as genotype and read depth data for samples at each genomic position. Read depth is a measure of sequencing coverage at each variant position for each patient. VCF files include lines of meta-information, a header containing sample IDs, and data rows for specific variant genomic locations. Since next generation sequencing is becoming increasingly accessible to researchers and clinicians, the ability to easily retrieve and visualize genomic data from VCF files is needed. This is beneficial for translational and personalized medicine.

Interpreting data from VCF files presents several challenges. The ability to process VCF files is limited by computational resources as the file size is often very large. To facilitate memory efficient data retrieval, existing VCF file parsing and visualization tools require users to preprocess their VCF files. This entails compressing and sorting VCF files by genomic position before either subsetting the file with an external program, such as VCFTools^1^, or indexing the files with Tabix^2^. Further, the VCF data structure is dense and difficult to interpret in its raw data format and requires data querying to draw insights. To facilitate efficient interpretation and data sharing, the need exists for user-friendly VCF file parsing and visualization tools.

We introduce “Visualization of Variants” (VIVA), a command line utility and Jupyter Notebook^3^ based tool for evaluating and sharing genomic data for variant analysis and quality control of sequencing experiments from VCF files. VIVA delivers flexibility, efficiency, and ease of use compared with similar, existing tools including vcfR^4^, IGV^5^, Genome Browser^6^, Genome Savant^7^, svviz^8^, and jvarkit – JfxNgs^9^. The distinguishing features of VIVA include: (1) No need for VCF file preprocessing (including compression, sorting, or indexing); (2) Ability to sort data by and visualize sample metadata; (3) No coding is necessary; (4) Variety of publication quality output formats; (5) Interactive HTML5 output for real-time data exploration and sharing; (6) Exports heatmap data as text file matrices to analyze using other tools.

## 2. RESULTS

In order to achieve this, VIVA employs the Julia programming language, a high-level, high-performance, dynamic programming language for numerical computing^10^. This is the first tool of its kind written in the Julia programming language and able to be integrated into workflows with other tools hosted by BioJulia, the Julia language community for biologists and bioinformaticians.

Using VIVA involves three main steps which are illustrated in Figure 1:

(1) User submits input files and chooses filtering options, if any are needed;
(2) VIVA reads VCF file and processes the data;
(3) VIVA creates graphs and exports output files.

**Figure 1:**
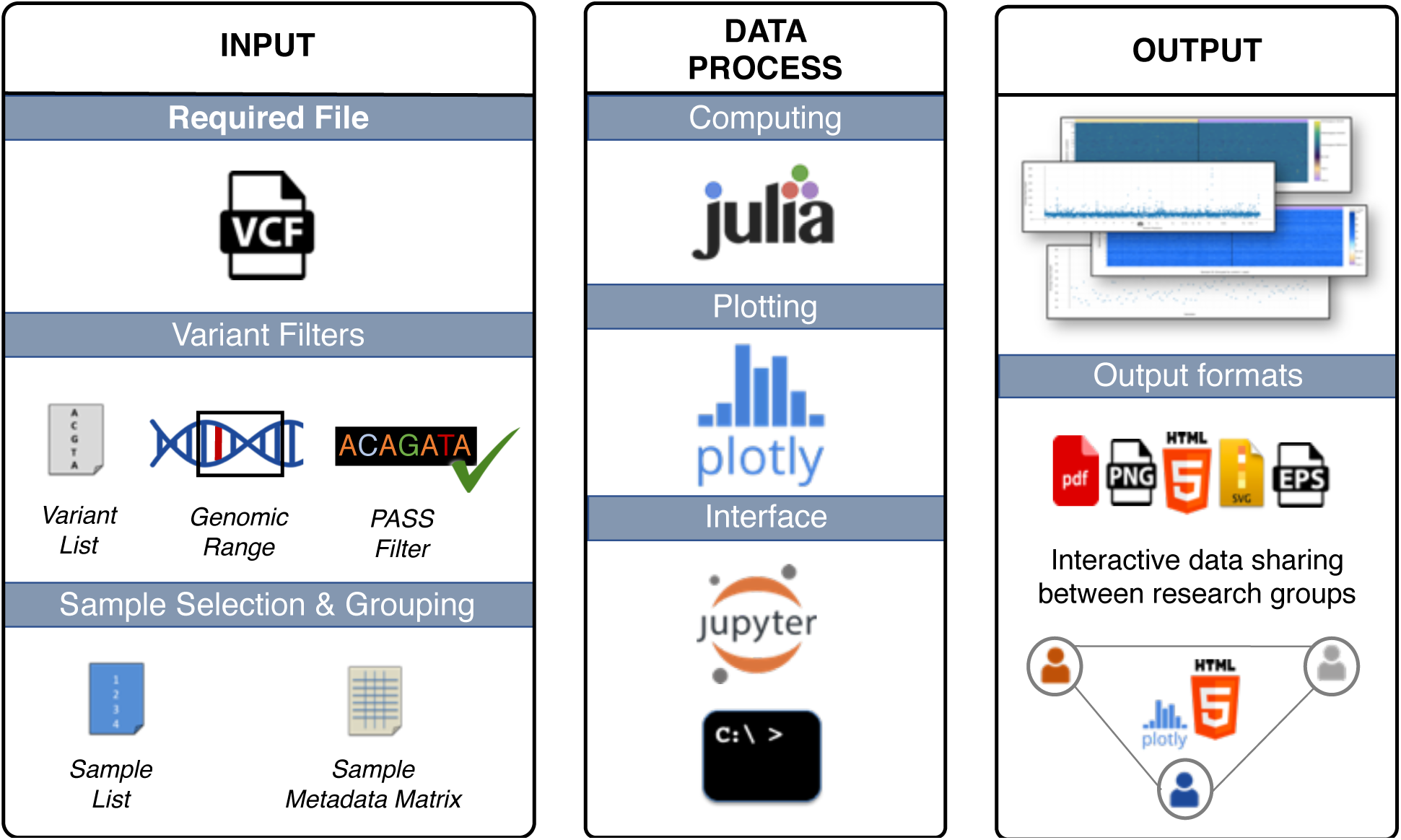
Workflow of VIVA. *INPUT*: VCF is a required file. Users can use one or any combination of variant filters, sample selection and grouping options. *DATA PROCESS*: Data processing requires Julia programing language and depends on several well-maintained Julia packages. Plotting uses the PlotlyJS.jl wrapper for Plotly. VIVA has two interface choices; users may use the program through a Jupyter Notebook or from the command line. *OUTPUT*: VIVA’s four visualization options include heatmaps of genotype and read depth data as well as scatter plots of average sample read depth and average variant read depth data. These visualizations can be saved in HTML, PDF, SVG, or EPS formats. HTML format enables users to share and analyze the data interactively between research groups which supports collaborative work environments.

Input data accepts four file formats. Examples of these input files are found in the documentation hosted at https://github.com/compbiocore/VariantVisualization.jl and are described in the Methods Section. The VCF file is the only required file.

There are three optional text file inputs for variant filtering and sample selection:

(1) *Variant List*: A list of specific variant positions of interest to include in visualizations. Users prepare a comma separated list in .csv format where the first column includes chromosome number and the second column includes genomic position;
(2) *Sample Metadata Matrix*: Users label their samples in a .csv file with phenotypic or experimental metadata information so the program can group samples with common traits and add this information to a heatmap of the genotype or read depth data. Any number of binary phenotypic traits or experimental conditions can be added to the matrix;
(3) *Sample List*: Users can select specific samples of interest to include in visualizations by submitting a .csv file containing sample IDs.

Since the number of data points allowed for visualization is limited both by the user’s computational resources available for plotting and pixels needed for display, we recommend using one or a combination of VIVA’s variant filtering options. These filtering options include:

(1) *Pass Filter*: Selects variant records that have passed filters selected during VCF generation;
(2) *Variant List Filter*: Uses the Variant List input file described above to select variant records that match a list of genomic positions; (3) *Genomic Range* Filter: Selects variants that lie within a given genomic range (Ex: chr1:8,900,000-12,000,000).

VIVA combines fast variant record filtering with visualization and data summarization without requiring allocation of a large amount of memory. This allows users with standard amounts of computational resources to analyze their whole VCF files. We have developed filtering functions with a low-memory-footprint by evaluating each line of the VCF file while only saving to memory variant lines that match filters. The VIVA command line and Jupyter Notebook tools depend on several Julia packages including VariantVisualization.jl, GeneticVariation.jl, ArgParse.jl, DataFrames.jl, PlotlyJS.jl. The use of these packages is explained in detail in the Methods section.

VIVA supports multiple visualization options. These include heatmaps of genotype or read depth data with samples in columns and variant positions in rows. Genotype heatmaps are categorical heatmaps that display the genotype values: homozygous reference, heterozygous variant, homozygous variant, or no call for all selected samples and variants. Read depth heatmaps are plots of continuous read depth values from 0-100, or no call. *“*No call” indicates that there was poor data quality during VCF generation to this point. Read depth outlier values greater than 100 are capped to avoid loss of resolution of low read depth values in visualizations. The read depth ceiling of 100 was chosen because, for most purposes, a read depth value of 30+ is adequate for inclusion in variant analysis^11,12^. VIVA can also generate scatter plots of average read depth across samples or variant positions. Users can use these to identify issues with sample preparation and variants that lie in difficult-to-sequence regions as demonstrated in Figure 2. Additionally, users can choose to save labeled data matrices of representative genotype values or continuous read depth values for analysis in external programs.

**Figure 2:**
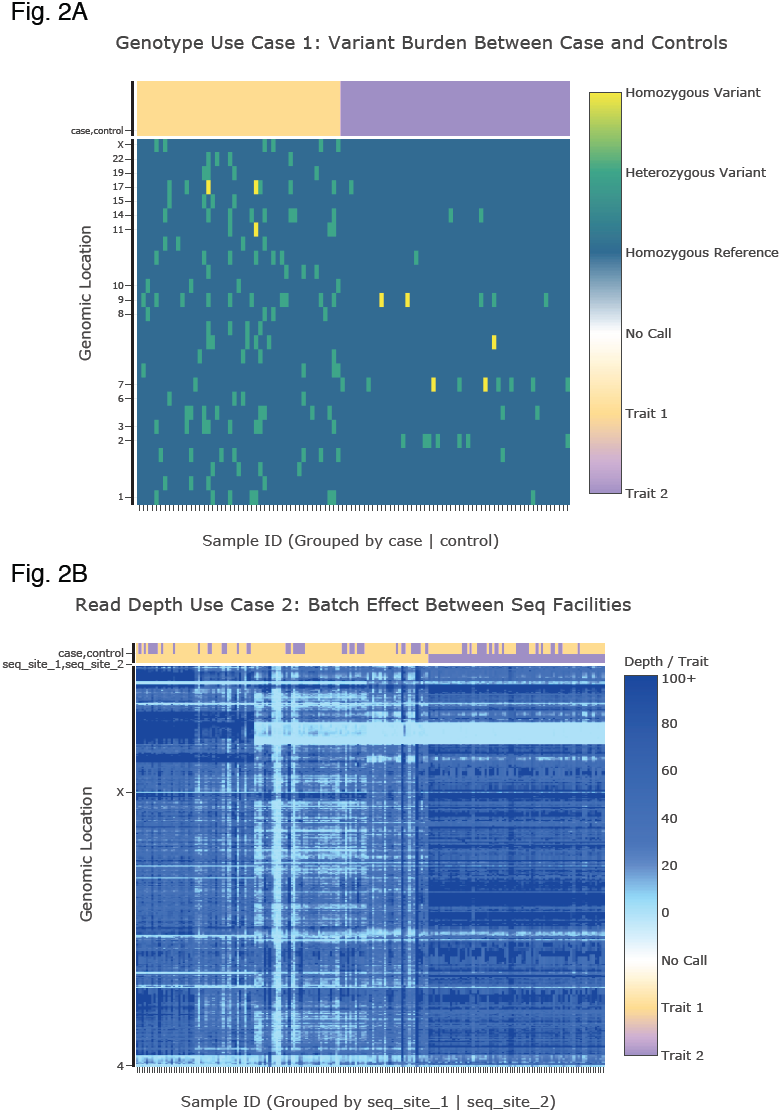
VIVA Use Cases. We present two use cases for VIVA. In both, unique variant positions are stored in rows and individual samples are stored in columns. We selected VIVA options to display only chromosome labels, rather than each specific chromosomal position, to create a cleaner presentation. In the first use case (**Fig. 2A**), we visualize a differential burden of putative disease associated variants between cases and controls by visualizing genotype data and grouping samples by case and control. In the second use case (**Fig. 2B**), we identify batch effects between samples sequenced at two separate facilities for a variant analysis study by visualizing read depth information and grouping samples by sequencing facility. We chose to visualize variants within chromosomes 4 and X arbitrarily to reduce the dimension of the data for memory efficient plotting.

VIVA exports plots in a variety of interactive and static graphic formats. HTML5 output facilitates interactive visualization and supports cursor hover-text, zooming, scrolling, and screen capture features. Cursor hover-text contains the sample ID, genomic position, and value for each data point in heatmaps and scatter plots. This format can be used for data sharing purposes and serves as a real-time data investigation tool. Static visualizations can be saved in PDF, SVG, PNG, and EPS formats for print and web publications and for presentations.

We believe that it is important to build a bioinformatics tool that can be used easily by non-programmers. While users with coding skills can benefit from and contribute to the open source code and command line tool, we have also implemented a Jupyter notebook-based web application. It is an interactive web tool that enables the combination of software code and visualization outputs in the same medium, with explanatory text supported by links to documentation.

Another important feature of VIVA is that it operates as a stand-alone application. Users do not need to submit their VCF files into the *cloud* or other remote environments which protects privacy. They can install VIVA locally or on a high-performance computing cluster.

VIVA is an open source tool built upon our Julia package, VariantVisualization.jl, and is freely available at https://github.com/compbiocore/VariantVisualization.jl. In order to run VIVA, users need to install the Julia Programming Language v1.1.0 onto their operating systems. The tool undergoes routine cross-platform testing by the continuous integration service, Travis CI, which is used for testing projects on GitHub. By building VIVA using an open source Julia package, we have made it easy for other developers to integrate VIVA into their programs and contribute to our code.

We evaluated VIVA’s performance with a test data set on a MacBook Pro with 2.9 GHz Intel Core i5 CPU running macOS High Sierra with 8 GB 1867 MHz DDR3. Our test data set was a 13.58 GB VCF file from a whole exome sequencing study containing 6,699,236 variants for 100 samples (human). We ran VIVA and selected 8700 variants-of-interest from our test VCF file with default options to generate all plots. We saved four outputs, including a scatter plot of average sample read depth, a scatter plot of average variant read depth, and heatmaps of genotype and read depth values, all in HTML file format. We ran 5 replicates of this test and found it took an average of 4 minutes and 13 seconds with a range of 2 seconds.

## 3. DISCUSSION

Grouping samples by metadata categories is broadly useful for comparative evaluation between samples such as identifying batch effect between groups of samples or differential presence of variants between sample groups in one VCF file. When sample grouping is implemented by the user, metadata is visualized in a subplot of colorbars at the top of heatmap visualizations. Users can add as many metadata category rows as they like, and can group by any one trait at a time. The ability to group by combinations of traits is under development for a future VIVA version.

VIVA’s variety of visualization options creates many use cases for high-throughput sequencing experiment quality control and variant analysis. We present two such use cases in Figure 2.

## 4. METHODS

### 4.1 Tool Architecture

VIVA exists as both a command line tool and as a Jupyter Notebook hosted utility. Both of these tools are built with VariantVisualization.jl, our Julia programming language package for VCF file parsing, data manipulation, and plotting. VariantVisualization.jl depends upon a variety of other Julia packages which are described in this Methods section. Both the command line tool and the Jupyter Notebook utility utilize functions from VariantVisualization.jl in the following sequence: variant record selection, collection of genotype or read depth values for selected variants into a numerical array, reordering the columns of the numerical array using sample metadata, selection of specific samples, and finally, plotting the resulting data.

The VIVA command line utility is called from the command line by calling the Julia language, the tool name, the name of the VCF file to visualize, and finally all usage options (‘julia viva -f file.vcf [options]’). VIVA options are evaluated and passed to VariantVisualization.jl functions within the command line utility by the ArgParse.jl Julia package. Users should reference the documentation hosted on the VariantVisualization.jl GitHub repository (https://github.com/compbiocore/VariantVisualization.jl) or run ‘julia viva –help’ for a list of all features and use instructions. User instructions for the Jupyter Notebook tool are contained within the Jupyter Notebook.

### 4.2 Data Input

The only required input file is the VCF file. VIVA specifically supports VCF files for human genomes and will not recognize chromosomes with non-standard names (outside of chr1-22, chrX/Y/M). VCF variant records do not need to be sorted by chromosome position. Chromosomes may be named in either ‘chr1…chrX’ or ‘1…X’ conventions which must be consistent across all chromosomes.

Three optional input files include:

(1) a list of specific variant positions to select;
(2) a sample metadata data table to group samples by;
(3) a list of samples to select.

The list of chromosome positions of interest must be formatted as a tab delimited .csv file with chromosome number in the first column and variant start position in the second column. An example of proper formatting can be found on the GitHub repository for VIVA under the path, tests/test_files/positions_list_test_4X_191. Chromosome number formatting should be consistent with chromosome number format in the VCF file (eg. either chr1 or 1). The second optional input file, a phenotypic data table, is a table saved in tab delimited .csv with information about each sample for grouping samples by common traits. There is an example of this file in the VIVA GitHub repository under tests/test_files/ sample_metadata_matrix .csv which can be used as a formatting guide. In this table, column names are sample ids, row names are comma separated group trait terms (‘case, control’), and cells contain a binary value for the sample/trait of interest. The third optional input file is a tab delimited list of sample ids to select for visualization and should be formatted to match the example file in the VIVA GitHub repository under tests/test_files/select_samples_list.txt.

### 4.3 Variant Record Filtering

Users set filtering and visualization options in the command line interface or in the Jupyter Notebook VIVA utility’s settings. Variant records are evaluated to match the filtering options and are stored in an array of chosen records. Selected records are converted into numerical arrays and sorted by chromosomes 1-22,X,Y,M for plotting. VIVA utilizes several Julia packages to read and process VCF files. VCFTools.jl is used to display the number of variant records and samples in the VCF file at the start of the program’s run. VIVA depends upon the GeneticVariation.jl Julia package to read data from VCF files. GeneticVariation.jl was chosen because it allows for easy parsing and data extraction from VCF records and is actively maintained by the BioJulia community. VCF files are read in the form of a VCF.Reader object which allows reading records one by one in an ∷IO stream to allow processing large VCF files without loading them into local memory. The VCF.Reader object holds all of the information contained in one row of a VCF file, including chromosome number, position, filter status, and genotype information for each patient.

Variant records can be selected using three optional filtering choices: “-- pass_filter”,”--list”, and “--range”. The first filtering option, “--pass_filter,” reads over all records and selects variants that pass QC filters chosen when producing the VCF file. If the record contains the string “PASS” in the “Filter” field, it is added into an array of records for visualization. The second filter option is “--list” and selects variant records from the user-provided list of chromosome start positions. The formatting of this list is specific and is described in the “Data Input” subsection of this Methods section. The VariantVisualization.jl package uses the io_sig_list_vcf_filter() function to iterate through each record in the VCF.Reader object and check if the record matches the list of user-defined chromosome positions. To save time, the function stops iterating through the records once the number of selected variant records matches the known number of records in the list of chromosome positions of interest. The third variant record filter option is “--range” and selects variant records with values in their chromosome and position fields that are within a user-specified chromosome range. This range must be within a single chromosome and defined in the specific format: “chr1:4000-50000000.”

If no filters are applied, large VCF files (with 100,000+ variants) will take a long time to process because they will have to be loaded into memory as an array of many variant records. The user must have enough RAM to load the VCF file into memory as an array of records. Generally this can be achieved on a shared computing cluster. As an important reminder, we do not recommend visualizing this many variants at a time. Heatmap visualizations are limited by pixel size, so visualized variants will lose definition at this scale.

### 4.4 Converting to Numerical Arrays

Once variant records have been selected, a numerical array is generated to contain either the genotype or read depth values for each variant in each sample. To visualize genotype values in a categorical heatmap, genotype values are converted into categorical representative values: no call = 0, homozygous reference = 1, heterozygous reference = 2, homozygous variant = 3. Chromosome number and positions are stored as row names and sample ids are stored as column names of these matrices. Column and row names are used for reordering and selecting samples as well as labeling plots. In addition to heatmap visualizations, users can generate scatter plots of average read depth values across samples as well as across variants. Users can identify problematic samples with low coverage by plotting average sample read depth. To do this, the means of read depth values for all selected variants are calculated for each sample and are plotted in a scatter plot. Similarly, users can identify hard to variant regions with low coverage by plotting average variant read depth. To plot average variant read depth, the mean of read depth values for each sample is calculated for each variant and are plotted in a scatter plot. Read depth outlier values are capped at 100 to scale the data for visualization, as is done with read depth heatmap plotting. For analysis in external programs, users can choose to save these numerical arrays as tables.

### 4.5 Sample Ordering and Selection

There are two options to manipulate the VCF data using sample ids. These options both depend upon the DataFrames.jl v0.11.7 package. Users can reorder the columns of the VCF file to explore trends across samples by supplying a matrix of sample metadata and sample ids. This is described in the “Data Input and Preprocessing” subsection of this Methods section.

To reorder sample columns, the numerical array of genotype or read depth data is converted into a DataFrame. Then the sample metadata matrix is grouped according to a chosen trait and the order of the sample ids contained within the sorted metadata matrix is used to reorder the.numerical DataFrame. To select columns, the numerical array is converted into a DataFrame and a new DataFrame is declared to only include columns with column names matching the sample ids provided in user defined list. These DataFrames are converted back into arrays for plotting.

### 4.6 Generating Plots

We have built our plotting functions using PlotlyJS.jl v0.10.2. PlotlyJS.jl is a Julia wrapper for plotly.js, an open-source JavaScript charting library. We used this library to build heatmap functions for plotting read depth and genotype data and to create summary scatter plots of average read depth values. We chose PlotlyJS.jl because it is very customizable, well maintained, and integrates with Rsvg.jl v0.2.1 to allow saving graphics in a variety of publication quality, scalable formats.

Numerical arrays of read depth or genotype values are plotted by a heatmap function to produce a categorical heatmap. Categorical genotype values 0, 1, 2, and 3 represent genotype conditions “no call”, “homozygous reference”, “heterozygous variant”, and “homozygous variant” and are plotted with the Viridis color palette. We chose this color palette by the recommendation of Nathaniel Smith and Stefan van der Walt who announced Viridis as the default colormap of the popular python plotting package, Matplotlib 2.0 at the SciPy 2015 Conference. They stated Viridis is accessible to viewers with color blindness, visually appealing, and able to be converted to grayscale^13^. Continuous read depth values are plotted in a continuous value heatmap using shades of blue that are reminiscent of ocean floor relief maps. This caps the maximum DP at 100 and prevents high read depth values from obscuring resolution of low read depth values which are usually of greater interest. They are optimized to show clear distinction between read depths in the range of 0-50 by coloring all read depth values over 100 the same, since read depth of greater than 30 is usually adequate for downstream analysis. Heatmap y-axes are labeled with chromosome positions using input matrix row names and x-axes are labeled with sample ids using column names of the input matrix. Save formats for all heatmap and scatter plots include PDF, HTML, SVG, PNG, and EPS. Interactive plots can be saved in HTML to be used for real-time data exploration and are easily shared with other researchers who don’t have VIVA installed. By default, VIVA saves graphics in HTML format. HTML plots are in HTML5 format and can be viewed in any browser and support zooming, panning, and hover labels over the cursor for real-time data exploration. Hover labels contain chromosome position and sample number labels. When HTML is chosen as the save format, the plot axes are not labeled with tick labels per sample id and per chromosome. Plots saved in any format other than HTML will have x and y-axis tick labels.

### 4.7 Jupyter Notebook

Jupyter is an open source computational notebook that combines code, descriptive text, and interactive output and has become the computational notebook of choice with data scientists. We have used the VariantVisualization.jl Julia package to set up a Jupyter Notebook with the full functionality of the command line tool to guide users unfamiliar with running bioinformatics tools from the command line through using VIVA. It includes a concise user manual in the first cell of the notebook. The next cells contain clearly labeled fields for entering the VCF file name and desired options. The user only needs to fill out the data input and option selection fields, then run the final cell to produce, save, and display interactive plots within the notebook. Users can re-run analysis with different settings more quickly in the notebook.

### 4.8 Software and Code Availability

The open source Julia command line tool, Juypter notebook, and Julia package are available at https://github.com/compbiocore/VariantVisualization.jl. Installation and comprehensive use instructions are detailed in the VIVA documentation which is available at https://github.com/compbiocore/VariantVisualization.jl. Julia package version numbers listed in this Methods section are subject to change as they are updated routinely as part of VIVA’s ongoing development, Instructions on reproducing figures in this article are detailed in the user manual as well.

## 5. CONCLUSIONS

In conclusion, we have built a visualization tool for exploratory analysis and generation of publication quality graphics for variant analysis projects using Variant Call Format (VCF) files. Researchers and clinicians can use VIVA to explore phenotypic and genotypic associations, batch effect on coverage, and differential incidence of variants between samples in their variant analysis experiments.

## ACKNOWLEDGEMENTS

We thank Ben J. Ward (The Clavijo Group, The Earlham Institute, Norwich Research Park), developer of GeneticVariation.jl, who provided frequent technical support while we developed our functions for memory efficient variant record extraction from large VCF files. We thank Spencer Lyon (Managing Director, Valorum Data), developer of PlotlyJS.jl, who provided technical support while we developed our plotting functions. We also thank the Center for Computation and Visualization (CCV) and the Computational Biology Core at Brown University for their support in testing VIVA’s performance on different system configurations. This work was supported by the National Institutes of Health (grants 5P20GM109035-04, 5P30GM114750) and the Kilguss Research Core at Women & Infants Hospital.

